# Antimicrobial peptides (AMPs) of lizards: a first comprehensive characterization of beta-defensins, ovo-defensins and cathelicidins from the Balearic lizard *Podarcis lilfordi* and closely related Lacertidae species

**DOI:** 10.1101/2025.02.20.639273

**Authors:** Katherin Otalora, Jessica Gomez-Garrido, Laura Baldo

## Abstract

Reptiles are remarkably resistant to infections, providing a critical system to understand diversity and evolution of the innate immune defense and its major players, the antimicrobial peptides (AMPs). Here we present the first comprehensive characterization of AMPs in the family Lacertidae with the objective of understanding their diversity and patterns of evolution.

By means of extensive genome mining, we first obtained a nearly complete catalogue of antimicrobial proteins from the Balearic lizard *Podarcis lilfordi*: 65 beta-defensins (BDs), eight ovo-defensins (OVODs), encompassing three proline-rich proteins (OVOD-PrAMPs), and four cathelicidins (CATHs). Using this fine-scale annotation we retrieved corresponding orthologues and closed paralogues from published Lacertidae species, *P. muralis, P. raffonei* and *Zootoca vivipara* (58 total AMPs). Comparative sequence analysis indicated that all AMPs consistently locate in chromosome 3 (BDs and OVODs) and chromosome 12 (CATHs), supporting a monophyletic origin of the reptilian antimicrobial defense. In *P. lilfordi*, the AMPs are arranged in clusters of mostly contiguous peptides, flanked by highly conserved marker proteins. All Lacertidae AMPs present a multiple exon structure (two to four) and a characteristic cysteine motif (six-cysteines in BDs, eight in OVODs and four in CATHs), consistently with previous findings in vertebrates. Comparative analyses support an ongoing process of gene expansion via duplication in tandem of both BDs and OVODs, whereas OVOD-PrAMPs and CATHs mostly present a one-to-one ortholog in all species. Despite this remarkable intra-genomic diversity, we also found multiple examples of distant species sharing identical or nearly identical peptides, providing clear evidence of convergent evolution.

Overall, these findings substantially increased our understanding of AMP diversity and evolution in reptiles and set the basis to explore adaptive polymorphism maintenance and mechanisms of antimicrobial defense.

## Introduction

Antimicrobial peptides (AMPs) are a large class of small peptides (15-150 amino acids), found abundantly in nature and playing a crucial role in the innate immune systems of all organisms, from bacteria to vertebrates (Lazzaro et al. 2020; Huan et al. 2020; Chakraborty et al. 2022; Erdem Büyükkiraz and Kesmen 2022). They exhibit an ample spectrum of antimicrobial effects, primarily directed against bacteria, viruses, fungi and parasites, but also a potent immunomodulatory activity (Chakraborty et al. 2022; Huan et al. 2020). They represent the first line of an organismal defense against pathogens, while also acting as critical modulators of the host-microbe interactions (Cardoso et al. 2022).

AMPs are typically derived from post-translational modification involving a proteolytic cleavage of large precursor proteins (Zhang and Gallo 2016). Most active peptides present a net positive charge and an amphipathic nature that allow direct membrane binding and disruption in target microbes (Zhang and Gallo 2016; Erdem Büyükkiraz and Kesmen 2022; Chakraborty et al. 2022). Anionic peptides have been also reported, although mechanisms of actions remain mostly unclear (Harris et al. 2009).

Coevolution of AMPs and pathogen resistance has led to a highly functionally and structurally diverse repertoire of AMPs (Huan et al. 2020; Erdem Büyükkiraz and Kesmen 2022) resulting from extensive gene duplication, pseudogenization, deletion as well as strong positive selection (Semple et al. 2005; Lazzaro et al. 2020). Nonetheless, evidence of polymorphism maintenance by balancing selection and convergent evolution has also been documented (Chakraborty et al. 2022; Lazzaro et al. 2020) indicating an overall complex pattern of molecular evolution. Hence, despite their ample spectrum of microbial targets, recent studies have shown that AMPs response can be remarkably specific, with single polymorphisms significantly influencing resistance to infections (Lazzaro et al. 2020; Hanson 2024).

Among the plethora of AMP-producing proteins found in vertebrates, the main classes are represented by defensins and cathelicidins (Huan et al. 2020). Both are characterized by presence of cysteine residues (four to eight) arranged in conserved motifs and forming disulfide bonds that contribute to their structural stability. Number of cysteine residues and spacing of cysteine-pair alignment currently define known subclasses (Semple et al. 2006; Semple and Dorin 2012). Beta-defensins are the most abundant and diverse subclass found in all vertebrates, known for their immunomodulatory and toxic activity, as well as their involvement in developmental processes, fertility, wound healing and cancer, among others (Semple and Dorin 2012). They produce small active peptides (20–50 residues) with a characteristic six-cysteine motif (Semple and Dorin 2012) and are typically arranged in large discrete clusters within a chromosome (Zhu & Gao 2013; Tu et al. 2015). Within the same syntenic block of the beta-defensins are found the ovo-defensins, a small subclass of AMP-producing proteins structurally similar to beta-defensins (containing a six-to-eight cysteine motif) so far described only in avian and reptile species, and named after their expression in the oviduct and putative role in egg protection (Whenham et al. 2015; Gong et al. 2010).

Cathelicidins are a small class of largely studied vertebrate-specific proteins, with multifunctional activities marked by both potent antimicrobial and immunomodulatory properties (Kościuczuk et al. 2012; Bals and Wilson 2003). They contain a typically four-cysteine motif and are synthesized as large precursor protein containing a conserved cathelin-like domain (CLD) and a functional and highly variable antimicrobial domain (Kościuczuk et al. 2012; Tomasinsig and Zanetti 2005).

While AMPs have been extensively studied in humans and other mammals, a comprehensive genomic characterization of AMP diversity is still lacking for most other vertebrate groups. Reptiles are an increasingly attractive model of antimicrobial defense studies as they represent the first branch evolving an innate immune response in extant vertebrates and are known for their remarkable resistance to infections (Field et al. 2022; Buchmann 2014); yet AMP studies in reptiles are still at their beginnings (Santana et al. 2021; Siddiqui et al. 2022).

Research over the past two decades has identified genes associated with the major classes of antimicrobial peptides in only few representative species of the four major reptilian orders (i.e. Squamata, Testudines, Rhynchocephalia and Crocodylia) (Santana et al. 2021). Within Squamata (snakes and lizards), AMP diversity have been documented so far in the Komodo dragon (*Varanus komodoensis*) (van Hoek et al. 2019), the green anole lizard (*Anolis carolinensis*) (Dalla Valle et al. 2012), and few snake species, e.g. the king cobra *Ophiophagus hannah* (Zhao et al. 2008) and the Burmese python *Python bivittatus* (Kim et al. 2017). No formal AMP characterization is yet available for any members of the family Lacertidae, including wall lizards (genus *Podarcis*).

Recently, the complete genome assembly and annotation of the Balearic wall lizard *Podarcis lilfordi* (Squamata, Lacertidae) has been published (Gomez-Garrido et al. 2023). This endemic species inhabits multiple islets surrounding the islands of Menorca, Mallorca and the Cabrera Island, where it has greatly diversified in morphology, pigmentation and life history traits (Pérez-Cembranos et al. 2020; Rotger et al. 2020). Due to the insularity of this species and the long-term monitoring of several of these populations (Rotger et al. 2016), the Balearic lizard is a growing natural model system for integrative studies of host and associated microbial symbionts (Otalora et al. 2024; Baldo et al. 2018, 2023) as well as pathogen resistance (unpublished) in a context of a rapidly changing environment.

Here we present the first and comprehensive identification of major classes of AMPs in the *P. lilfordi* genome by means of extensive genomic search. In an effort to understand AMP diversity and evolution within Lacertidae, we additionally mined currently published Lacertidae genomes including the common wall lizard, *P. muralis* (Andrade et al. 2019), the Aeolian wall lizard, *P. raffonei* (Gabrielli et al. 2023) and the viviparous lizard, *Z. vivipara* (Yurchenko et al. 2020), providing a first overlook at the antimicrobial defense repertoire for this reptilian family. Finally, comparative analyses of newly discovered AMPs along with previously published data from the Komodo dragon (family Varanidae) (van Hoek et al. 2019) and the green anole lizard (family Dactyloidae) (Dalla Valle et al. 2012) allowed us to explore major patterns of AMP diversity and evolution in reptiles.

## Results

### AMP genomic clusters in *P. lilfordi*

We performed an extensive screening of the *P. lilfordi* genome, applying the known full-length beta-defensins from Komodo dragon and the green anole, and refining the search according to the presence of a characteristic cysteine motif and specific flanking marker genes. We were able to identify two major AMP cluster areas in chromosome 3 and 12 of the *P. lilfordi* genome (Figure 1).

**Figure 1.**
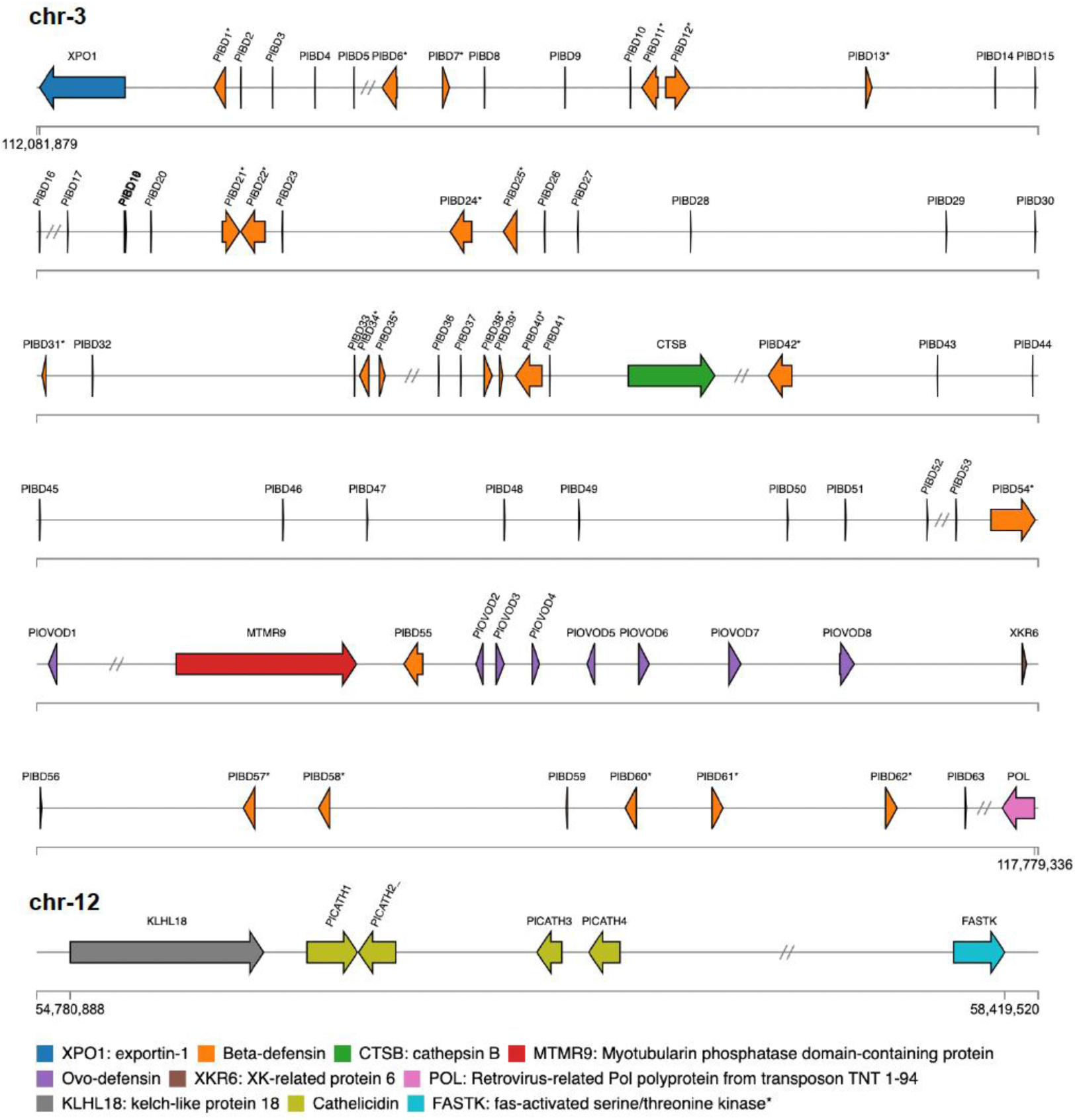
*P. lilfordi* genome localization of beta-defensins (chr-3), ovo-defensins (chr-3) and cathelicidins (chr-12). Beta-defensins include both partial and complete ORFs (marked with an asterisk). Marker genes for major clusters are also shown. Long intergenic regions are shortened and marked by a double //.

Chr-3 cluster (112164644-117711852 bp, about 5.5 Mb) includes 71 putative defensins: 63 beta-defensins (BDs) and 8 ovo-defensins (OVODs). The defensin block is flanked by the exportin-1 protein (XPO1) and the retrovirus-related Pol polyprotein from transposon TNT 1-94 (POL), with the cathepsin B protein (CTSB) falling within the block.

Ovo-defensins are found within the beta-defensin block, flanked by the myotubularin phosphatase domain-containing protein 9 (MTMR9) and the XK, Kell blood group complex subunit-related family, member 6 protein (XKR6).

Chr-12 cluster (54832594-58419520 bp, about 3.5 Mb) includes four cathelicidins (CATHs) arranged in a single block, flanked by the kelch-like family member 18 protein (KLHL18) and the fas-activated serine/threonine kinase protein (FASTK) (Figure 1).

### Characterization of *P. lilfordi* beta-defensins

All 63 BDs are arranged into two distinct clusters within chr-3, separated by ovo-defensins: one encompassing PlBD1-PlBD55 and the other PlBD56-63, separated by more than 2,000,000 bases (Figure 1 and Supplemental Table S1). All BDs identified present a six-cysteine motif (29 and 35 amino acid (AA)) with the following conserved spacing: C-X(6-7)-C-X(3-5)-C-X(8-14)-C-X(5-7)-CC (Figure 2). In addition, most sequences show two highly conserved glycine (G) residues at positions 7 and 28, and an isoleucine (I) at position 27 of the alignment (Figure 2). Both PlBD12 and PlBD22 carry a double cysteine motif in tandem at the C-terminal, summing a total of 65 AMPs.

**Figure 2.**
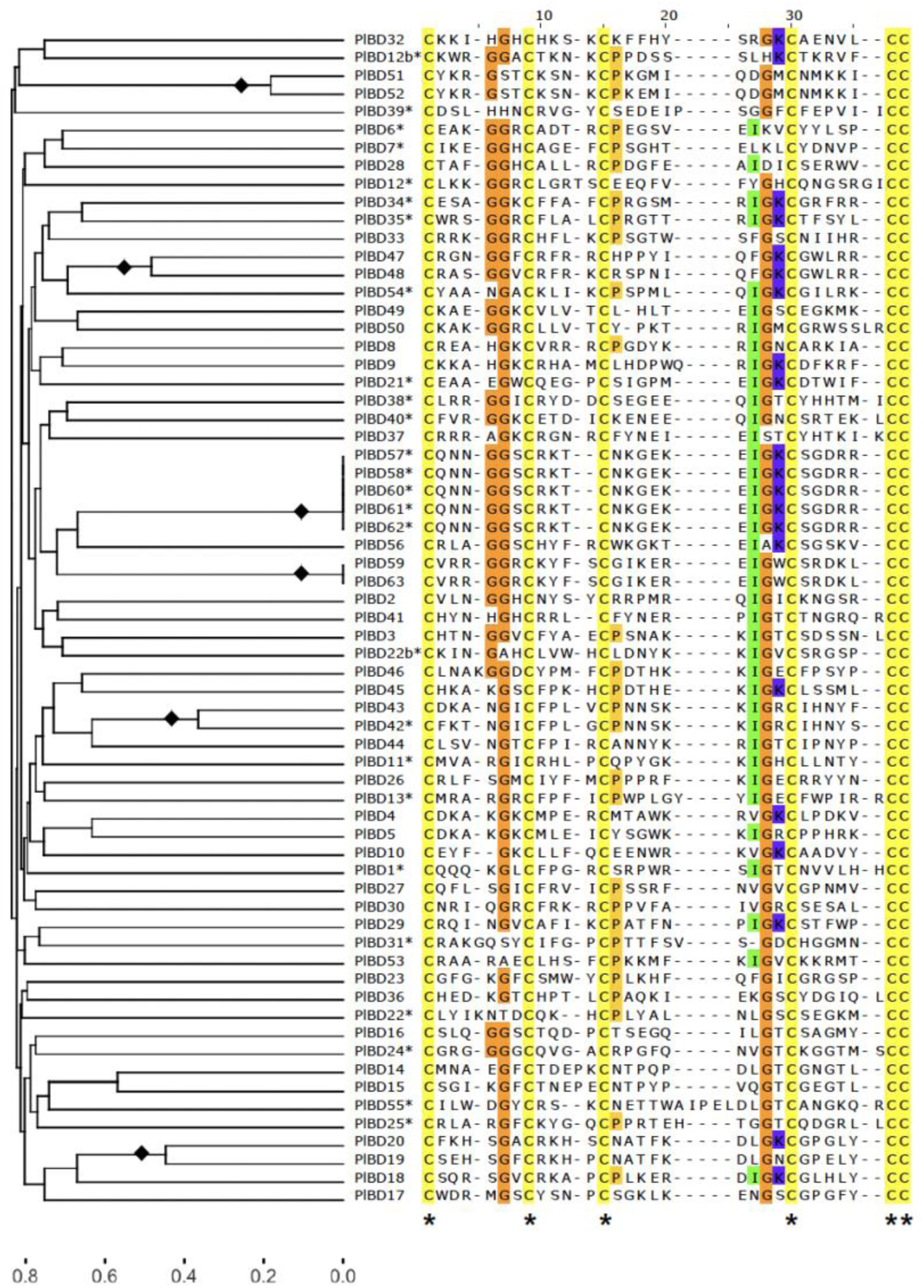
Alignment and UPGMA dendrogram of *P. lilfordi* putative beta-defensin peptides (N=65) showing the six-cysteine domain (C-X(6-7)-C-X(3-5)-C-X(8-14)-C-X(5-7)-CC). Alignment colors show residues above 30% identity threshold. Shared cysteine residues are marked with an asterisk. Dendrogram height corresponds to the square root of sequence identity distances. Branch squares highlight closely related peptides (cutoff at 0.5 height, corresponding to <25% of dissimilarity). An asterisk in sequence IDs correspond to peptides with complete ORFs. PlBD12 and PlBD22 carried a double motif, and the second motif was labelled with “b”.

Most peptides present a cationic charge, with isoelectric points ranging between 6.8 to 10.7, with only 13 out of the 65 peptides showing a negative or no charge (Supplemental Table S1). According to protein homology searches, all peptides (at least 30 AA) have their best hits to an antimicrobial protein/beta-defensin, with confidence >80% and sequence coverage >70%, except for PlBD39 (Supplemental Table S2). Prediction of secondary structure indicates that most cysteine domains containing two to three beta-sheets.

The UPGMA (unweighted pair group method with arithmetic mean) dendrogram identified six clusters of similar sequences (<25% dissimilarity based on an arbitrary cutoff), typically grouping contiguous or spatially proximal peptides within the chromosome (e.g., PlBD51-52, PlBD47-48, PlBD19-20, see also Figure 1). Two of these clusters encompass multiple BDs with identical motifs (PlBD57-58,60-62 and PlBD59,63), suggesting recent duplications in tandem.

Of the 65 beta-defensin motifs identified, we were able to detect the complete open reading frame (ORF) for 24 of them (Figure 3). According to published *P. lilfordi* transcriptomic data from multiple individuals and tissues (Bioproject: PRJNA897120, but also see genome browser https://denovo.cnag.cat/genomes/podarcis/browse/rPodLil1.2_annot/), fourteen of these BDs were transcribed, guiding our gene annotation (Supplemental Table S1).

**Figure 3.**
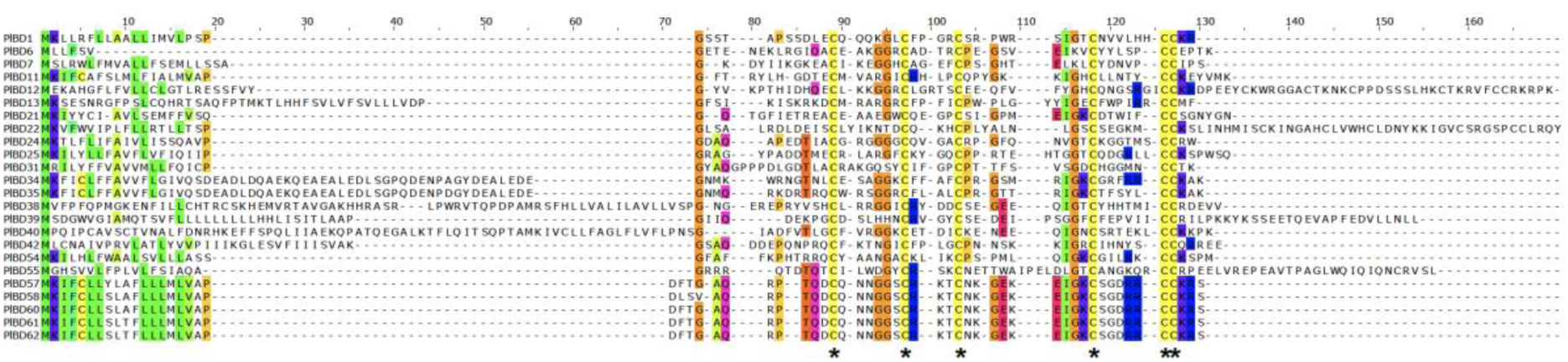
Alignment of *P. lilfordi* complete beta-defensin proteins (N= 24). Proteins encompass two to four exons (see Supplemental Table S1 for intron breaks). The six-cysteine residues are marked with an asterisk. Alignment colors show residues above 30% identity threshold. PlBD12 and 22 carry a double cysteine motif (the second motif was not aligned).

Complete protein size ranges from 53 to 118 AA arranged from two to four exons, with the majority presenting two-three exons, with large exon size variation (exon-1: 6-55 AA, exon-2: 21-69 AA, exon-3: 0-49 AA, exon-4: 0-45 AA) (see Supplemental Table S1 for intron position). They contain a N-terminal with a putative signal peptide (first exon and part of the second exon), followed by the cysteine motif at the C-terminal encoding for the AMP. Structural homology searches based on complete proteins confirm strong similarity to previously identified beta-defensins, with confidence >90%, although with generally low sequence identity (35-65%) (Supplemental Table S2). Again, only PlBD39 shows poor confidence (<80%).

While full proteins carry an overall substantial AA diversity at both N and C-terminal, the same cluster of BDs that showed full identity at the cysteine motif (PlBD58-62, Figure 2) also display near AA identity at full protein length (Figure 3), therefore supporting recent whole-gene duplication.

### Comparative analysis of beta-defensins across reptiles

To understand patterns of beta-defensin diversity and evolution within Lacertidae, we performed extensive BLAST searches that enabled to retrieve several complete proteins from closely related species to *P. lilfordi* (hereafter named Pl): 11 from *P. raffonei* (hereafter Pr, eight newly identified), seven from *P. muralis* (Pm, six new) and nine from *Z. vivipara* (Zv, six new). Most newly characterized beta-defensins were previously annotated as uncharacterized ncRNA or mRNA (see Supplemental Table S3 for AccNos).

All reptiles share the following six-cysteine motif: C-X(6-7)-C-X(3-7)-C-X(8-14)-C-X(4-7)-CC (see Supplemental Figure S1 for alignment).

The Neighbor-Joining (NJ) tree (Figure 4A) identified fourteen multispecies clades (bootstrap support >70) and only one clade of *P. lilfordi*-specific sequences (clade 34).

**Figure 4.**
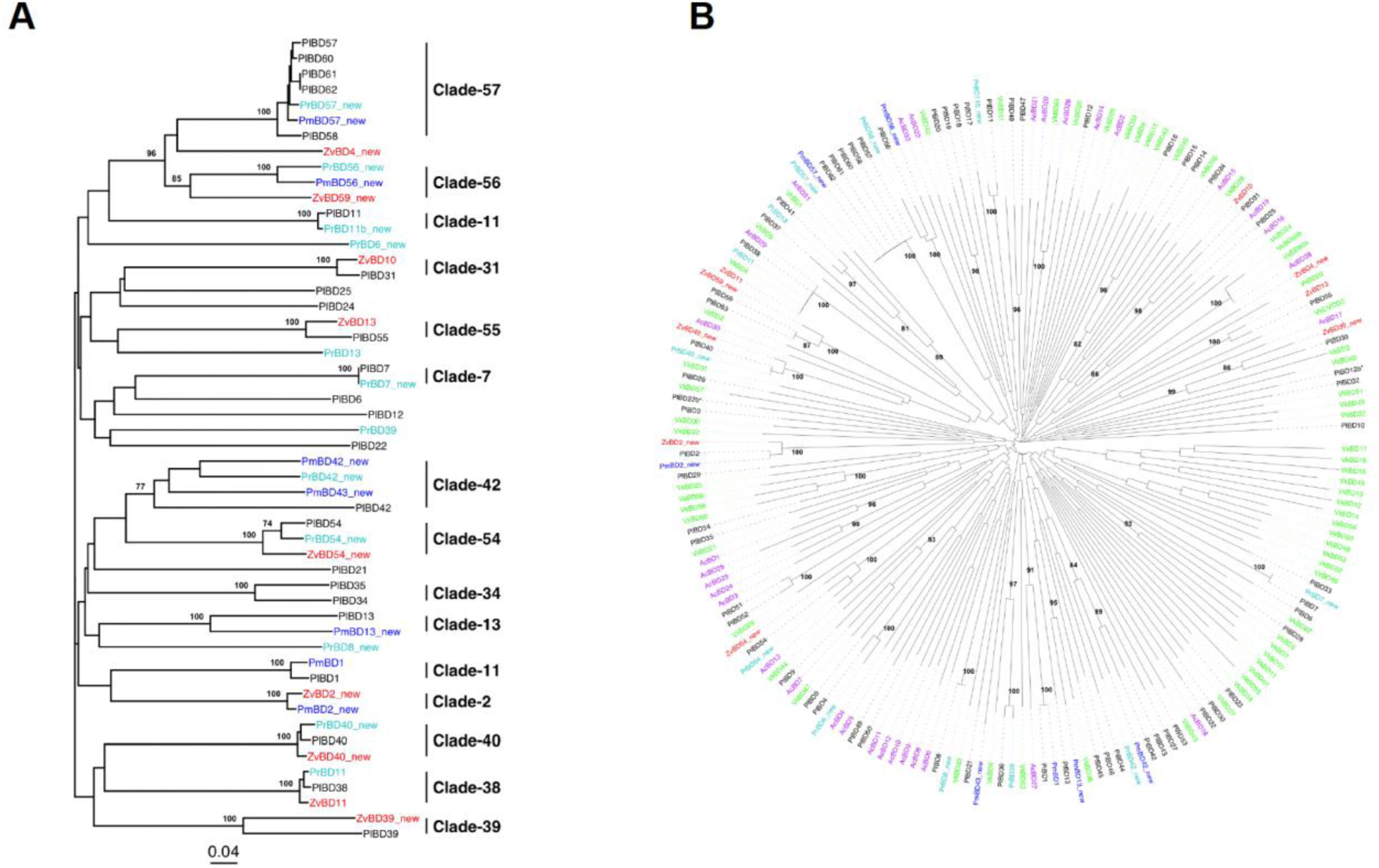
Beta-defensin comparative analysis across multiple related reptile species. A) NJ tree based on complete proteins (N= 47). Major clusters (bootstrap support>70) are highlighted and named according to the PlBD reference within the clade. B) NJ tree based on the shared cysteine domain from all five species (C-X(6-7)-C-X(3-7)-C-X(8-14)-C-X(4-7)-CC) (N =190 sequences), highlighting major clusters (bootstrap support>80). Proteins were named according to their public annotation; if uncharacterized, they were numbered according to their closest *P. lilfordi* sequence and marked with ‘new’ (see Supplemental Table S3 for species AccNos). Pl: *P. lilfordi*; Pm: *P. muralis*, Pr: *P. raffonei*; Zv: *Z. vivipara*; Vk: *V. komodoensis*; Ac: *A. carolinensis*.

Clades 54, 56 and 57 depict species relative distances according to their expected taxonomic grouping, showing all *Podarcis* spp. clustering together, with Zv at a basal position, supporting orthology. Few clades show high protein similarity among species, although without species or genus resolution: these include clade-38 and 40 (grouping *Podarcis* spp. and Zv), clade-1 (Pm-Pl), clade-2 (Pm-Zv) and clade-7 and 11 (Pl-Pr) (Figure 4A). Interestingly, clade-7 shows full identity of BD7 between Pr and Pl, suggesting putative convergent evolution (but see Discussion).

Due to a potential incomplete sampling of the full repertoire of beta-defensin full proteins from *Podarcis* spp. and Zv, we further explored pattern of BD evolution in reptiles based on cysteine motifs only, using a larger dataset encompassing also the Komodo dragon and the green anole lizard (N=190 sequences: Pl=65, Pm = 7, Pr= 12, Zv = 9, *A. carolinensis* (Ac) = 32, and *V. komodoensis* (Vk)= 65) (Supplemental Table S3).

The NJ tree (Figure 4B) highlights three main aspects: within species recent duplications, species-specific lineages, and across species peptide convergence. For recent duplications, aside from the several examples found in Pl (commented above), we did not detect any other recent intraspecific duplication for the other species (except the previously reported VkBD80a and b). Moreover, most lineages appear to be species-specific. Nonetheless, both patterns could also result from pervasive gene loss and/or incomplete sampling of beta-defensin diversity in all six species under study, unclear at present. The most notable aspect is the presence of eight clusters (bootstrap value>79) with motif identity across species (Figure 4B), with instances of motif identity is found for all species pairwise, although more frequently within the *Podarcis* genus (Pl-Pr: five clades; Pm-Pl: two clades; Ac-Vk: two clades, Pl-Pr-Zv-Ac-Vk: one clade). Notably, the clade encompassing AcBD29, PlBD38, VkBD4, ZvBD11 and PrBD11 (corresponding to clade-38 in Figure 4A) shows motif identity among five of the six species under study (excluding Pm), with representatives of the three distinct reptilian families (Lacertidae, Varanidae and Dactyloidae).

### Characterization of *P. lilfordi* ovo-defensins

We detected a total of eight complete ovo-defensin proteins in chr-3 of *P. lilfordi*, tightly clustered within or near the region flanked by MTMR9 and XKR6 proteins (Figure 1 and Supplemental Table S4). Protein size ranges from 64 to 239 AA arranged in two exons: the first exon putatively encoding the signal peptides (about 21 AA), whereas the second exon includes the putative AMP (Figure 5).

**Figure 5.**
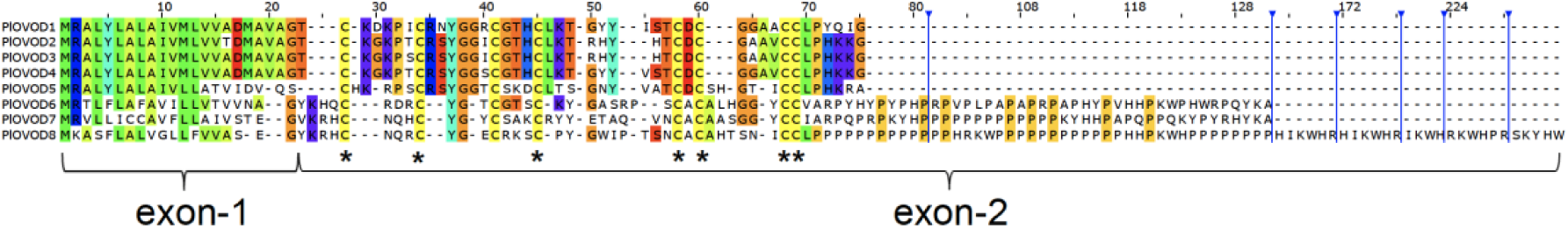
Alignment of *P. lilfordi* complete ovo-defensin proteins (N= 8). All proteins share a two-exons structure with a common intron position and the eight-cysteine motif C-X(3-5)-C-X(3-6)-C-X3-C-X(8-9)-C-X1-C-X(4-6)-CC, with a positive net charge. Blue lines mark the start of long stretches of prolines (>10) at OVOD7 and OVOD8. Shared cysteine residues are highlighted with an asterisk.

Intron size is relatively small, varying between 1.3 kb and 2.3 kb, except for OVOD1, where the intron is substantially larger (3.5 kb). All OVODs show an eight-cysteine motif with a highly conserved spacing: C-X(3-5)-C-X(3-6)-C-X3-C-X(8-9)-C-X1-C-X(4-6)-CC. PlOVOD7 presents two additional CC in the signal peptide. All peptides show a conserved Tyrosine (Y) and a Glycine (G) at position 37 and 38, respectively (Figure 5). PlOVOD6-8 are proline-rich proteins (PrAMPs), with PlOVOD7 and 8 presenting long stretches of up to 31 consecutive proline (i.e., proline repeats) (see Figure 5 for position of proline repeats and Supplemental Table S4 for complete sequences). In OVOD7 and 8, proline repeats are interspaced by repetitive lysine-tryptophan-histidine (KWH) and lysine-tyrosine-histidine (KYH) motifs. Two of the OVODs were found to be expressed according to transcriptome data (OVOD1 and OVOD4) (Supplemental Table S4).

All OVODs present a highly cationic charge (3.5 to 20) at exon 2 (i.e. the putative AMP), with isoelectric points ranging between 8.1 and 10.5 (Supplemental Table S4). Protein homology search indicated a poor confidence assignment (<71%) and coverage (<58%) for most peptides (Supplemental Table S2). Best hit is given for PlOVOD5 to a hydrolase/hydrolase inhibitor (carboxypeptidase inhibitor) (96.8% of confidence and 57% of coverage), and PlOVOD1 to an antimicrobial peptide (82% and 52%) (Supplemental Table S2). The remaining proteins do not present any significant match (>75% of confidence), likely due to the poor classification currently available for this group of defensins and to their intrinsically disordered structure.

OVOD2-4 share a large similarity (90-97% of AA identity), with PlOVOD2 and PlOVOD3 differing by a single AA (Figure 5) as a likely result of a recent duplication event. OVODs that are proximal in the chromosome tend to share higher protein similarity (Figure 1), in line with a process of duplication in tandem, as also observed for BDs. OVOD1-5 share little homology with OVOD6-8 (i.e., OVOD-PrAMPs, 7-24% of AA identity between them); aside from a conserved eight-cysteine residues motif, a two-exon structure, and a closed location in chr-3 (Figure 1), these two groups of OVODs might represent distinct protein families or subfamilies.

### Comparative analysis of ovo-defensins across reptiles

By means of extensive BLAST searches and genome mining we were able to retrieve several complete PlOVOD orthologs/paralogs from closely related species, none of which was previously identified: five from Zv, four from Pm, and eight from Pr (Supplemental Table S3). To these sequences, we added three previously described partial OVODs from Vk that met the minimum requirement of eight-cysteine residues (i.e., VkOVOD3, 4 and 6) Given the high fragmentation of current genome draft for *Varanus*, we were unable to retrieve the full proteins for this species.

The overall eight-cysteine motif across all reptiles was highly conserved, with a constant spacing between the third and fourth cysteine, and between the fifth and the sixth cysteine: C-X(3-6)-C-X(3-6)-C-X3-C-X(8-12)-C-X1-C-X(4-6)-CC (Supplemental Figure S2). Notably, OVOD6-8 represent proline-rich proteins in all Lacertidae, with OVOD7-8 showing long stretches of prolines interspaced by the conserved motif repeats KW(H) and KYH in all species (for Pm, only in OVOD8) (Supplemental Table S3 and Figure S2).

The NJ tree based on complete proteins identified five main clades (A-D, bootstrap support>80) (Figure 6).

**Figure 6:**
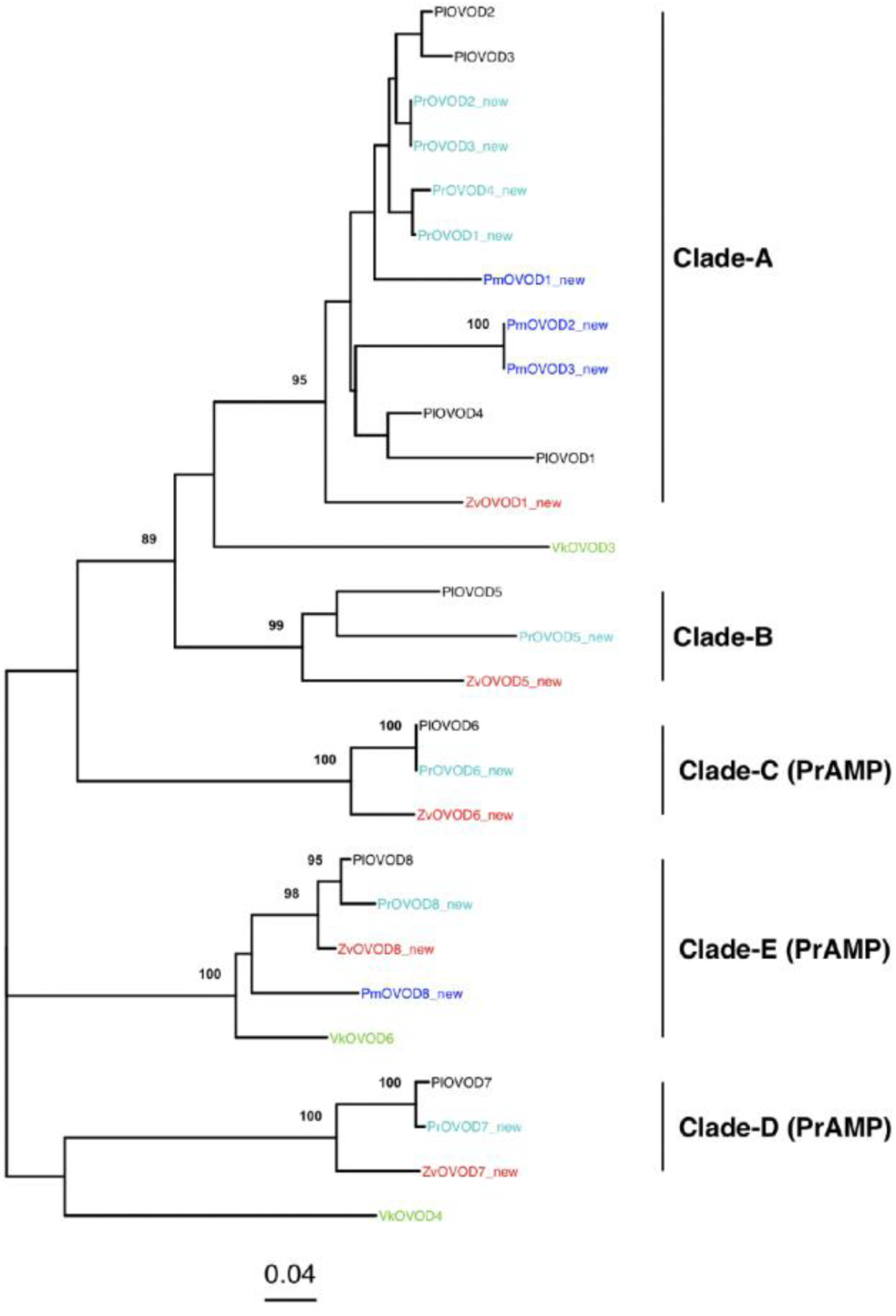
NJ tree based on complete ovo-defensin proteins (partial only for Vk) from multiple related reptile species (N=17). All species shared the following cysteine motif: C-X(3-6)-C-X(3-6)-C-X3-C-X(8-12)-C-X1-C-X(4-6)-CC. Major clades (bootstrap support>80) are labelled as A-E. Recent gene duplications are framed. New proteins were marked with ‘new’ and numbered according to their closest Pl sequence for Zv or to their sequential order in the chromosome for Pr and Pm (see Supplemental Table S3). Pl: *P. lilfordi*; Pm: *P. muralis*, Pr: *P. raffonei*; Zv: *Z. vivipara*; Vk: *V. komodoensis*.

Within Clade A, we can observe several instances of recent intraspecific duplication within all *Podarcis* spp., including Pl (OVOD2-3, with one AA difference, and OVOD1-4), Pr (OVOD2-3, OVOD1-4, each pair sharing identical cysteine motifs) and Pm (OVOD2-3 with full protein identity). Clade A and B group together with high support, suggesting a common origin by past lineage duplication. The remaining three clades C-E include the proline-rich proteins OVOD6-8, all showing a one-to-one ortholog in all Lacertidae, with no recent duplications. OVOD8 shows cysteine motif identity (although not full identity at whole gene level) across Pl, Pr, Pm and Zv, and OVOD6 between Pl and Pr (Supplemental Figure S2).

### Characterization of *P. lilfordi* cathelicidins

We identified four complete cathelicidin proteins in chr-12 of *P. lilfordi* (PlCHAT1-4), flanked by KLHL18 and FASTK proteins (Figure 1 and Supplemental Table S5). PlCATH2 was labelled as PlCATH2-OH following the annotation of a closed related protein in Zv, named as ZvCATH2-OH to maintain previous annotation from the king cobra *Ophiophagus hannah* species, where this protein was first described (Zhao et al. 2008). PlCATH1 and PlCATH2-OH are spaced by only 266 bp, although found in opposite strands (Supplemental Table S5).

Protein size of the four CATHs vary between 161 to 171 AA and are all arranged in four exons (Figure 7). All four CATHs were expressed according to transcriptome data. They exhibit a conserved four-cysteines domain with a highly conserved spacing: C-X10-C-X10-C-X(16-19)-C.

**Figure 7:**
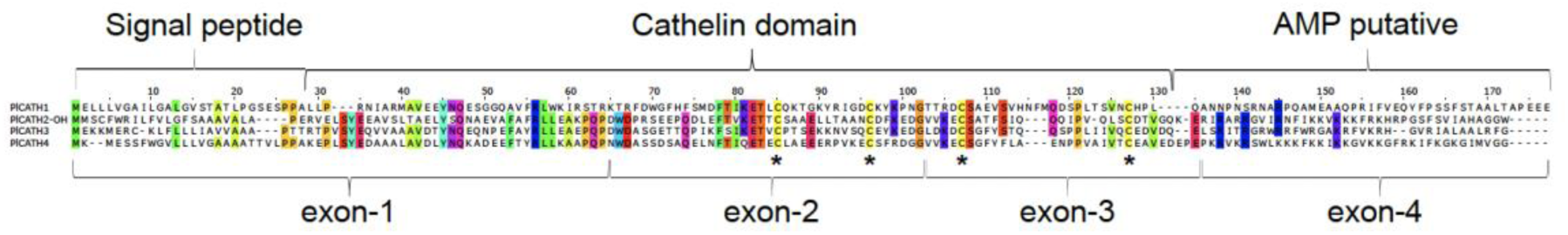
Alignment of *P. lilfordi* complete cathelicidin proteins (inactive precursor) (N= 4). All proteins share a four-exons structure and the four-cysteine motif C-X10-C-X10-C-X(16-19)-C. Main domain structure (signal peptide, cathelin domain and putative AMP) was predicted by homology searches. Shared cysteine residues are highlighted with an asterisk.

Homology searches returned best matches to porcine CATHs, with 99.9% of confidence and coverage above 50% for all four proteins (Supplemental Table S2). Folding predictions indicate that all four PlCATHs represent inactive precursors, structured in three domains: 1) a variable N-terminal domain with the signal peptide of slightly variable length (first 23-27 AA), 2) a conserved “cathelin-like domain” (CLD) containing the cysteine motif (100-101 AA long), with one alpha helix and four beta-sheets, and 3) a highly variable C-terminal domain containing the putative functional AMP (36-42 AA), with proteolytic cleavage predicted at position 130 (Figure 7).

Except for PlCATH1, all other AMP domains presented a highly cationic charge (11.5-15) due to multiple Lys (K) and Arg (R) residues, with isoelectric points ranging between 12 to 13 (Supplemental Table S5). AMP domain from PlCHAT1 largely differed in AA content as lacking K and G residues and presents a net negative charge (-2).

Overall protein similarity among the four CATHs is low (<40%), indicating no recent duplications.

### Comparative analysis of cathelicidins across reptiles

For comparative purposes, we retrieved all available CATHs from the Lacertidae under study as well as identified six novel ones: five from Pm (one newly identified), four from Pr (three new), and four from Zv (two new). We additionally included three CATHs from Vk and Ac, all previously described. Most novel CATHs were annotated as uncharacterized mRNA (see Supplemental Table S3 for AccNos).

During sequence alignment, we found that a single CATH-coding mRNA sequence from Pm, Pr and Zv included two consecutive cathelicidin proteins: a first protein with high similarity to PlCATH4, followed by a second protein with high similarity to PlCATH3. Coexpression of CATH4-CATH3 was not observed in Pl, where the two proteins are separated by 5901bp (Figure 1 and Supplemental Table S5). For alignment purposes, we manually split the mRNA to recover individual proteins and aligned them along with the other sequences.

The multispecies alignment indicates that all reptiles under study share a highly conserved four-cysteine motif: C-X(9-10)-C-X10-C-X(17-21)-C (Supplemental Figure S3). The Valine-130 (following the last cysteine residue), predicted as the proteolytic cleavage for the active peptide in Pl, was also conserved across all reptiles in CATH2-4, but not in CATH1 (Supplemental Figure S3).

The NJ tree based on full proteins (Figure 8) clearly depicts four highly supported clusters of one-to-one orthologous proteins (Clade 1-4), largely recovering the main phylogenetic position of the two Lacertidae genera: all *Podarcis* species together (with no further resolution at species level), with Zv at a basal node.

**Figure 8:**
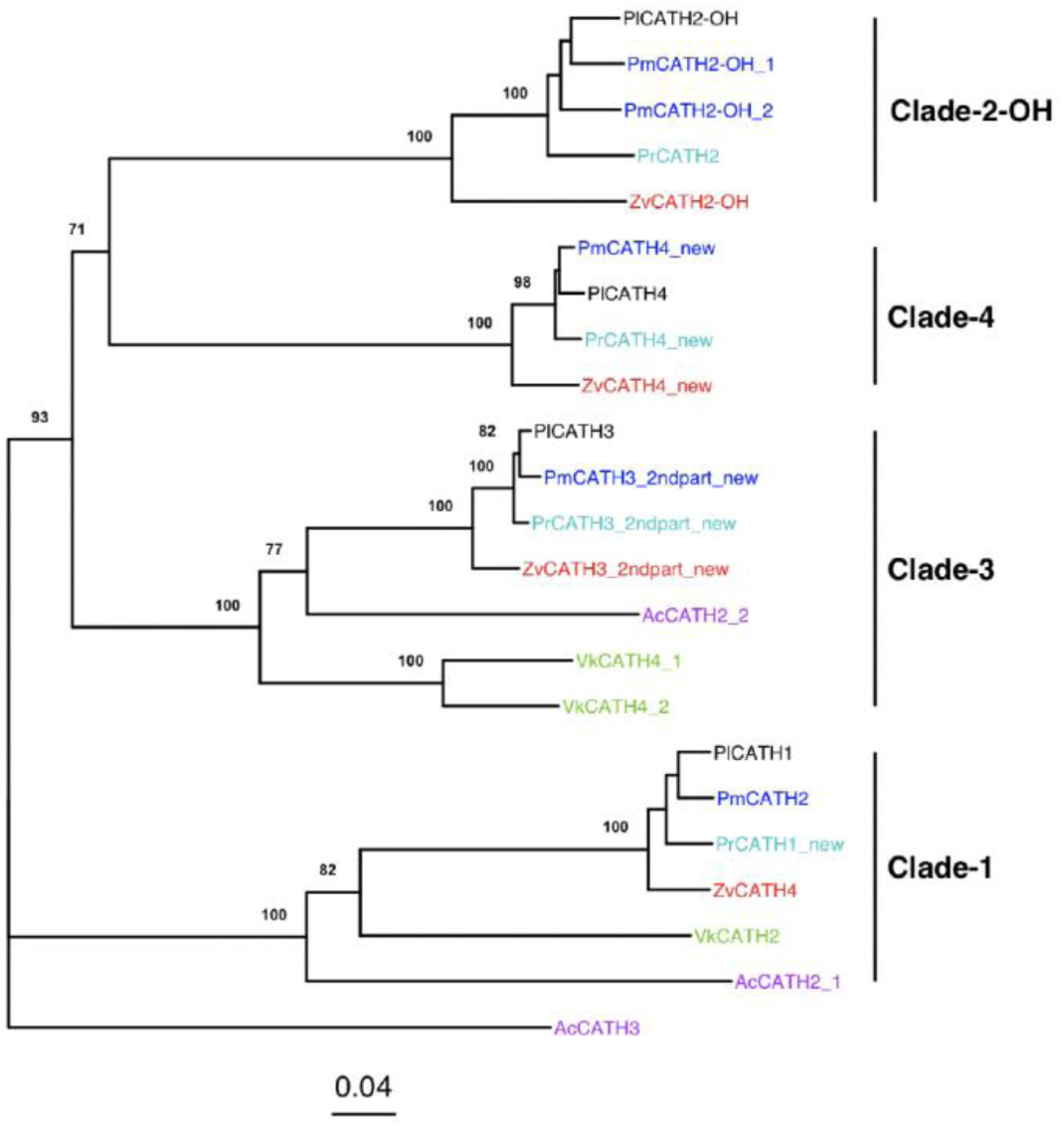
NJ tree based on complete cathelicidin proteins from multiple related reptile species (N=23). All reptiles shared the four-cysteine motif C-X(9-10)-C-X10-C-X(17-21)-C. Major clades (bootstrap support>70) are labelled based on the *P. lilfordi* reference protein.

Moreover, the species Ac and Vk are basal to both clade 1 and 3, with Vk presenting two closely related paralogs in CATH3. Both species do not present any orthologue to clade 2-OH or clade 4, although, given the incompleteness of *V. komodoensis* genome, this absence could be also ascribed to failure in sequence identification for this species. A recent CATH duplication is only observed in Pm, with two proteins falling within the clade 2-OH.

## Discussion

The innate immune system of lizards remains a vastly understudied field of research (Field et al. 2022; van Hoek 2014). In this study we provided the first comprehensive genomic characterization of major AMP families in the Balearic lizard *P. lilfordi* and closely related species within the family Lacertidae.

For all Squamata species under study (including the Komodo dragon and the green anole), AMPs consistently clustered in the same two chromosomes, chr-3 for beta- and ovo-defensins, and chr-12 for cathelicidins, making these regions important repositories of the reptile antimicrobial defense. The same genomic organization has been previously reported for crocodiles and birds, but not for mammals (Santana et al. 2021), in line with an avian and reptilian monophyletic origin. Within these regions, *P. lilfordi* flanking proteins of AMP clusters (i.e., XPO1 and CTSB for beta-defensins, XKR6 and MTMR9 for ovo-defensins, and FASTK and KHLL18 for cathelicidins) were consistent with those previously described in other reptiles, but also mammals (Santana et al. 2021; van Hoek 2014; Dalla Valle et al. 2012) underscoring their importance as a critical reference markers for AMP cluster identification in reptiles.

### Beta-defensins diversity and evolution in reptiles

Beta-defensins represent the most expanded and diverse AMP family in vertebrates (Tu et al. 2015). No current studies are available for Lacertidae, with Komodo dragon and the green anole being the closest species presenting a comprehensive characterization of beta-defensins (van Hoek et al. 2019; Dalla Valle et al. 2012).

Diversity in *P. lilfordi* (65 peptides, 24 with complete ORFs) largely aligned with that previously described in the green anole (N=32) (Dalla Valle et al. 2012) and the Komodo dragon (N=65) (van Hoek et al. 2019). All proteins presented substantial diversity in the amino acid sequence, length, molecular weight, and isoelectric points; yet they were all characterized by the canonical six-cysteine motif and a mostly cationic charge reported for all vertebrates (Tu et al. 2015; Semple and Dorin 2012; Nava et al. 2009). Moreover, complete beta-defensins exhibited a broad structural variation, with two to four exons (two being the most recurrent), that also conforms to that observed in the green anole (Dalla Valle et al. 2012) and the Komodo dragon (van Hoek et al. 2019).

In *P. lilfordi* we observed that sequence similarity was typically higher among chromosomally adjacent peptides or full proteins, suggesting a process of paralogous expansion via duplication in tandem. Instances of identity or virtual identity among pairs of contiguous peptides further indicate that this process is still ongoing. These observations are in line with previous research supporting a major role of gene duplication in the overall increasing diversity of this gene family (Lazzaro et al. 2020; Semple and Dorin 2012; Tu et al. 2015). Notably, we also found instances of beta-defensins (PlBD12 and 22) carrying two consecutive cysteine motifs at the C-terminal, a structural organization that was previously observed in other reptiles, but not in mammals (Guyot et al. 2022), and that suggests the concomitant expression of multiple AMPs. Given the high sequence dissimilarity between the two consecutive domains, these double motifs are unlikely to be derived from intra-protein duplication in tandem; rather they most likely resulted from a sequence translocation or other chromosomal rearrangements still to clarify.

In line with this complex pattern of intra-genomic diversification, comparative beta-defensin analyses across reptiles recovered a generally low phylogenetic signal due to pervasive gene duplication, but also putative gene loss and rapid diversification at the active peptide (Tennessen 2005). We note that, while the extensive AMP genome screening of *P. lifordi* provided a nearly complete catalogue of beta-defensins for this species, current AMP characterization in the other Lacertidae genomes might be incomplete, limiting our current inferences on the relative role of gene loss and specie-specific gene duplication in the observed diversity. Nonetheless, we found several clades of putative true orthologs across all Lacertidae (Figure 4A), supporting a substantial level of evolutionary conservation for some of the beta-defensins, as previously observed in mammals (Nava et al. 2009; Lazzaro et al. 2020). These clades were characterized by variable branch lengths, suggesting heterogeneous rates of protein changes along species-specific lineages, likely driven by protein niche functional specialization in host-pathogen interactions (Semple et al. 2006; Chakraborty et al. 2022). Interestingly, we found several instances of cysteine motif identity or high similarity between members of the genus *Podarcis*, particularly between the two closest related species *P. lilfordi* and *P. raffonei* (Yang et al. 2022), but also among members of distant genera/families (e.g., BD38 was shared among the three families under study, Lacertidae, Dactyloidae and Varanidae). This pattern strongly supports a major role of convergent evolution, rather than strong stabilizing selection, in shaping beta-defensin diversity. High rates of AMP convergent evolution have been previously observed in both vertebrates (including humans) and invertebrates (mainly *Drosophila*) (Unckless and Lazzaro 2016), where distant taxa were shown to share identical alleles or specific polymorphisms as a result of convergent adaptive immunity (Lazzaro et al. 2020). For instance, AcBD29 from *A. carolinensis* displayed the same AMP motif across all five species under study (Figure 4B); in the green anole, this anionic peptide is known to be expressed in all tissues, although its function remains unknown (Dalla Valle et al. 2012). Another peptide, AcBD27, which formed a well-supported clade with *P. lilfordi* (PlBD1) and *P. muralis* (PmBD1), is known to be highly expressed in wounded lizards, particularly following tail loss, where it helps preventing infection despite the absence of inflammation, a key factor in successful regeneration (Alibardi 2013b, 2013a). While species-specific studies are clearly needed to clarify a putative functional equivalence of the same beta-defensin peptide across species, the above examples can set the basis for exploring putative conserved mechanisms of action.

### Ovo-defensins and PrAMPs diversity and evolution in Lacertidae

Unlike beta-defensins, ovo-defensins represents a smaller subfamily of still largely understudied defensins, currently described only in birds and few reptiles (but none in Lacertidae), where they appear to be specifically expressed in the oviduct and contribute to the egg immune defense (Whenham et al. 2015; Zhang et al. 2019; van Hoek et al. 2019).

The eight OVODs identified in *P. lilfordi* were all located in the same chr-3 cluster region, flanked by conserved marker genes, MTMR9 and XKR6, previously described for other vertebrates (Zhang et al. 2019), indicating a strong monophyletic origin for this subfamily, although with few chromosomal rearrangements (i.e. PlOVOD1 was located immediately outside the gene cluster). We confirmed the two-exon structure and a higher divergence at exon-2 previously reported for avian and reptilian ovo-defensins (Whenham et al. 2015). Compared to beta-defensins, OVODs present two extra cysteines (eight total), a finding that is in contrast with previous reports of a six-cysteine motif for OVODs in reptiles (Whenham et al. 2015). Notably, this eight cys-motif and two-exon structure was observed in all reptiles under study (both Lacertidae species and the Komodo dragon), strongly suggesting they represent specific structural features of this protein subfamily. Differences in cysteine number and arrangement between beta- and ovo-defensins may underscore fundamental functional differences that are currently unclear (Erdem Büyükkiraz and Kesmen 2022). Despite structural commonalities and a proximal chromosomal arrangement, the eight PlOVODs encompassed a remarkable amino acid diversity and pattern of evolution that might warrant a further categorization into distinct subfamilies of defensins. While ovo-defensins from clade A appear to have undergone an extensive process of gene expansion by duplication in tandem in all *Podarcis* species (Figure 6), the remaining clades (B-E) present a single ortholog for each species, supporting their relative evolutionary stability. Interestingly, these clades encompasses proline-rich AMPs (PrAMPs) (Lai et al. 2019) containing either multiple non-adjacent prolines (P) (OVOD6) or proline repeats at the C-terminal, typically interspaced by motifs rich in lysine (K), tryptophan (W), histidine (H), arginine (R) and tyrosine (Y) (OVOD7 and 8) (Figure 5). Their structure and lack of any significant protein folding suggest they might represent “extended AMPs”, a class of AMPs known to contain high proportions of P or W residues and no regular secondary structure elements (Nguyen et al. 2011). Notably, both P and W-rich peptides have been shown to carry potent antimicrobial activity for their ability to permeate the outer membrane and additionally target intracellular pathways (Lai et al. 2019; Mishra et al. 2018; Hernandez-Gordillo et al. 2013). For instance, recent studies have demonstrated that the polyproline host defense peptide bactenecin 7 (Bac7) can bind the wall polysaccharide biofilm of hypermucoviscous bacteria, such as *Klebsiella pneumoniae*, actively disrupting bacteria cell wall (De Los Santos et al. 2024; Fleeman and Davies 2022). Additionally, KW repeats (here observed in OVOD8) were shown to increase AMP bactericidal activity (Gopal et al. 2013), whereas KY residues (observed in OVOD7) are known to cooperate with H in facilitating DNA cleavage in DNase proteins (Korn et al. 2002).

Interestingly, we found several instances of cys-motif identity of OVOD-PrAMPs across species, with OVOD8 carrying the same motif in all Lacertidae. As observed for beta-defensins, this suggests a major role of convergent evolution in shaping PrAMPs diversity.

### Cathelicidins diversity and evolution in Lacertidae

Cathelicidins are a small and ancient family of potent AMPs, with large functional plasticity and a broad spectrum of actions against a wide range of microorganisms (Alford et al. 2020; Kościuczuk et al. 2012; Tossi et al. 2024). Presently, they have been described in several reptile species of the order Squamata (Tossi et al. 2024), including the Komodo dragon (van Hoek et al. 2019), with no formal description yet available for any Lacertidae species, except for gene annotation of few representative cathelicidin proteins from public genomes.

The four cathelicidin proteins identified in *P. lilfordi* (CATH1-4) were found to be tightly clustered in chromosome 12, flanked by the same conserved genes, FASTK and KLHL18, previously reported for birds and reptiles (Cheng et al. 2015; van Hoek et al. 2019; Tossi et al. 2024). They all exhibit a classical four-exon precursor structure of all vertebrates characterized by a variable signal peptide, a conserved four-cysteine motif and a highly divergent C-terminal carrying the active peptide (Izadpanah and Gallo 2005; Gennaro and Zanetti 2000; Zhong et al. 2020). Of the four cathelicidins, three present a mature peptide with a highly cationic net charge (>10), in line with the average positive charge found for reptiles, including the Komodo dragon (Alford et al. 2020). Notably, CATH1 was the only protein presenting an anionic AMP in all reptiles under study (net charge -1 to -2) and an overall AA composition largely distinct from the other cathelicidins, suggesting a rather distinctive mechanism of action for this protein. To date a single anionic cathelicidin has been formally described and functionally characterized in salamanders, where it is produced by the skin and carries a potent anti-inflammatory and wound healing activity, although not direct antimicrobial effect (Luo et al. 2021). Functional studies are required to assess a potential role of this cathelicidin in lizard wound healing after injuries or tail autotomy, a conserved anti-depredatory mechanism of all lizards (Alibardi 2014).

All Lacertidae presented four cathelicidins (with one exception in *P. muralis*), all located in chr-12, with a one-to-one ortholog in all species, supporting a monophyletic origin and phylogenetic conservation along this taxonomic lineage. Three of these orthologues clades (2-4) clustered into a major well-supported clade, including also representatives of the Komodo dragon and the green anole, pointing to a common origin by a past duplication within Squamata. A fifth CATH was found in *P. muralis* and resulted from a single recent duplication in Clade-2-OH, underlying that cathelicidin gene duplication is still an ongoing process, although less pervasive than observed in defensins (Tossi et al. 2024). Interestingly, we found that CATH3 and CATH4 were coexpressed into a single large RNA in all Lacertidae, except in *P. lilfordi*, suggesting a post-translational process to concomitantly generate the two distinct active peptides. We found no previous reports in literature of such an RNA arrangement for cathelicidins. Nonetheless, we observed similar instances of multidomain structure also in beta-defensins (see above), suggesting it might provide some functional gain, unclear at present (Guyot et al. 2022). In line with this, AMP coexpression models in experimental systems were recently shown to effectively enhance the antimicrobial activity and immunity (Chen et al. 2022; Kuddus et al. 2017), pointing to potential cooperative interactions among naturally coexpressed AMPs, which is worth further investigation.

### Conclusions

Our findings substantially broaden current knowledge on the diversity of AMPs in reptiles and their mechanisms of evolution, marked by gene duplications and fast evolution. The comprehensive characterization of major AMP families in the Balearic lizard *P. lilfordi* can now aid in the fine-scale annotation of these peptides in other reptiles, therefore facilitating comparative studies among Squamata species for a comprehensive understanding of gene duplication/loss within specific reptilian lineages. Particularly, the identified cysteine motif identity and high protein similarity across species provide the background to explore conserved mechanisms of antimicrobial action of these peptides in reptiles, as well as their value as candidates for new therapeutic developments.

As for the Balearic lizard *P. lilfordi*, identification of major players in innate immunity sets the basis to understand the role of AMPs in their ecoimmunology and potential response of these insular populations against infection threats that are rapidly increasing with increased levels of anthropization in these islands. Healthy *P. lilfordi* populations play significant roles in their ecosystems and a fine understanding of their genetic potential in immune response can offer new insights into resilience mechanisms, guiding species-specific conservation efforts for these isolated populations. Additionally, future comparative analysis across all extant *P. lilfordi* populations will provide an exciting opportunity for AMP population-level genetic studies aimed to shed light on the rapid evolution and adaptive polymorphism maintenance of the reptilian antimicrobial defense.

## Methods

### Identification of candidate defensin and cathelicidin genes in *P. lilfordi*

For the identification of candidate defensin and cathelicidin genes within the *P. lilfordi* genome we employed a multiple genomic approach.

First, we used the protein definitions file (PODLIA.protein_definitions.txt) available at https://denovo.cnag.cat/podarcis and retrieved the already annotated AMPs from the recently published genome (Gomez-Garrido et al. 2023). These included four putative beta-defensins, one cathelicidin, and no ovo-defensins.

Second, we identified the chromosome and cluster boundaries for defensins and cathelicidins by searching for presence of conserved flanking marker genes, as documented in literature (Dalla Valle et al. 2012; van Hoek et al. 2019; Cheng et al. 2015; van Hoek 2014). Specifically, for beta and ovo-defensins we targeted the exportin-1 protein (XPO1), the cathepsin B protein (CTSB), the XK, Kell blood group complex subunit-related family, member 6 (XKR6) and Myotubularin phosphatase domain-containing protein 9 (MTMR9) (van Hoek et al. 2019; Zhang et al. 2019). For cathelicidins we targeted the fas-activated serine/threonine kinase (FASTK) and kelch-like family member 18 (KLHL18) (Cheng et al. 2015; van Hoek et al. 2019).

Third, we retrieved available AMPs from the closest reptile species for which these peptides have been identified, the *V. komodoensis (*Komodo dragon) (van Hoek et al. 2019), and the *A. carolinensis* (the green anole) (Dalla Valle et al. 2012), and used them as queries in a TBLASTN search (Altschul et al. 1990) against the *P. lilfordi* genome. The Komodo dragon includes 65 described beta-defensins peptides, six ovo-defensins (only mature peptides), and three cathelicidins (complete genes) (van Hoek et al. 2019); the green lizard includes 32 beta-defensins (complete mRNA), three cathelicidins (complete genes) and no ovo-defensins (Dalla Valle et al. 2012). For each protein family, we applied relaxed TBLASTN search parameters (e-value = 1, gap open/extension = 8/2) to maximize output retrieval. The BLAST outputs were then filtered according cysteine (C) motifs: for beta-defensins, we filtered for presence of six cysteines and a relaxed cysteine spacing from van Hoek et al. (2019); for ovo-defensins, we retained best hits containing a terminal “CC” motif and between six to eight total cysteine residues (van Hoek et al. 2019; Dalla Valle et al. 2012); finally, for cathelicidins, we retained best hits with at least four cysteine residues (Tomasinsig and Zanetti 2005). Throughout this systematic approach, we verified and corrected the orientation of all sequences, prioritizing non-overlapping sequences with the highest bit score. All analyses were performed with custom scripts in R software (2023).

### Identification of Open-Reading-Frames (ORFs) and gene annotation

To obtain the complete Open-Reading-Frames (ORFs) for the identified AMPs that passed the above filters, we retrieved all available full mRNA and proteins from *A. carolinensis* (32 beta-defensins mRNA and four cathelicidin complete proteins) and V. *komodoensis* (only three cathelicidins proteins). All proteins were queried using the program exonerate v2.4.0 (Slater and Birney 2005) in protein2genome mode against chr-3 (defensins) and chr-12 (for cathelicidins) of the *P. lilfordi* genome. For ovo-defensins, as no complete proteins were available, partial ovo-defensin proteins newly identified from *P. lilfordi* were searched against the NCBI nt database: best RNA hits were translated to proteins and searched back against the *P. lilfordi* genome. Finally, the available *P. lilfordi* transcriptomic data (Illumina and PacBio) from multiple tissues and individuals (Bioproject: PRJNA897120) was used to correctly identify exons and introns. This approach allowed us to confirm most of the genes that had been previously identified using partial sequences as well as annotate novel genes.

For naming newly identified proteins/peptides in *P. lilfordi*, we used a standard code used for other species including the genus and species initials as prefix, followed by the protein code and a number corresponding to order in the genome location (Dalla Valle et al. 2012; van Hoek et al. 2019). New candidate proteins identified from Lacertidae were similarly named and numbered according to the *P. lilfordi* closest sequence, when available.

AMP chromosome location was visualized with the R function GC_chart in *geneviewer* R library. For the UPGMA (unweighted pair group method with arithmetic mean) dendrogram, a protein distance matrix was estimated with the R function ‘dist.alignment’ (matrix = “identity”, gap=T) in *seqinr* library, and a dendrogram generated using *hclust* with height corresponding to the square root of identity distances.

### Physical properties and 3D peptide prediction

Peptide physical properties were predicted by EMBOSS tool pepstats https://www.ebi.ac.uk/jdispatcher/seqstats/emboss_pepstats. Protein structure (domain definition, 2D and 3D) was predicted by homology searches with Phyre2 (Kelley et al. 2015).

### Comparative analysis across reptile species

To explore patterns of diversity and evolution of AMPs in Lacertidae, we used both available genome annotation and a combination of blast searches (BlastP and TBLASTN) to retrieve AMP sequences from three additional species from which genome assemblies were published: the Aeolian wall lizard, *P. raffonei* (Gabrielli et al. 2023); the common wall lizard, *P. muralis* (Andrade et al. 2019); and the viviparous lizard, *Z. vivipara* (Yurchenko et al. 2020). Additionally, we integrated available sequences from the Komodo dragon and the green lizard. Amino acid sequences were aligned using MUSCLE *vs*. 5.0 (Edgar 2004), visualized and manually curated in Jalview *vs*. 2.11.3.3 (Waterhouse et al. 2009). As homology inference within the same protein family is low due to the high rate of amino acid changes and extensive duplications, alignment curation was done following these general guides: a) we first aligned C-residues within the conserved motif, b) we maximized homology among highly similar sequences, c) we optimized similarity within the conserved regions among all sequences, and c) for highly variable regions (i.e. functional peptide), we minimized indels among divergent sequences.

To identify potential orthologs and paralogs among species, a Neighbor Joining (NJ) tree (Saitou and Nei 1987) was built with MEGA 4.0 (Tamura et al., 2007) with 1000 bootstrap values. The resulting trees were graphed with the *ggtree* R package (Yu et al. 2017).

## Data access

All data presented in this study was previously available. AccNos are referenced in the text and Supplemental Table S3.

## Fundings

This study was funded by MICIU/AEI/ 10.13039/501100011033 and by “ERDF A way of making Europe”, grant PID2022-141578NB-C22 to L.B. Doctoral support to K. O. was from the Doctoral Scholarship, Colombian Ministry of Science, Technology and Innovation (MINCIENCIAS885/2020 to K.O.). Institutional support to CNAG was from the Spanish Ministry of Science, Innovation and Universities, Fondo de Investigaciones Sanitarias cofunded with ERDF funds (PI19/01772), the 2014-2020 Smart Growth Operating Program, and the Generalitat de Catalunya through the Departament de Recerca i Universitats and Departament de Salut.

## Author contributions

L.B. conceived the study. L.B., K.O. and J. G. analyzed the results. L.B. and K.O. wrote the paper. All authors reviewed and contributed to the final version of the paper. Correspondence to L.B.

## Competing Interests Statement

The authors declare no conflict of interests.

## Supplementary Material

**Table S1**: Beta-defensin peptides and complete proteins from *P. lilfordi*.

**Table S2:** Summary of beta-defensins, ovo-defensins and cathelicidin protein homology results by Phyre2.

**Table S3:** List of AccNos of complete defensin and cathelicidin proteins from published reptile genomes. Newly identified are indicated with ‘new’.

**Table S4:** Ovo-defensin complete proteins from *P. lilfordi*.

**Table S5:** Cathelicidin proteins from *P. lilfordi*.

**Figure S1:** Multispecies alignment of beta-defensins proteins (partial) from the Lacertidae species under study (Pm, Pr, Pl and Zv).

**Figure S2:** Multispecies alignment of ovo-defensins proteins (complete) from the Lacertidae species under study and the Komodo dragon (Pm, Pr, Pl, Zv and Vk).

**Figure S3:** Multispecies alignment of cathelicidins proteins (partial) from Lacertidae species, the Komodo dragon and the green anole lizard (Pm, Pr, Pl, Zv, Vk and Ac).

## Notes

### Competing Interest Statement

The authors have declared no competing interest.

